# Antiseptic Agents Elicit Short-Term, Personalized and Body Site-Specific Shifts in Resident Skin Bacterial Communities

**DOI:** 10.1101/230193

**Authors:** Adam J. SanMiguel, Jacquelyn S. Meisel, Joseph Horwinski, Qi Zheng, Charles W. Bradley, Elizabeth A. Grice

## Abstract

Despite critical functions in cutaneous health and disease, it is unclear how resident skin microbial communities are altered by topical antimicrobial interventions commonly used in personal and clinical settings. Here we show that acute exposure to antiseptic treatments elicits rapid but short-term depletion of microbial community diversity and membership. Thirteen subjects were enrolled in a longitudinal treatment study to analyze the effects of topical treatments (ethanol, povidone-iodine, chlorhexidine, water) on the skin microbiome at two skin sites of disparate microenvironment: forearm and back. Treatment effects were highly dependent on personalized and body site-specific colonization signatures, which concealed community dynamics at the population level when not accounted for in this analysis. The magnitude of disruption was influenced by the identity and abundance of particular bacterial inhabitants. Lowly abundant members of the skin microbiota were more likely to be displaced, and subsequently replaced by the most abundant taxa prior to treatment. Members of the skin commensal family Propionibactericeae were particularly resilient to treatment, suggesting a distinct competitive advantage in the face of disturbance. These results provide insight into the stability and resilience of the skin microbiome, while establishing the impact of topical antiseptic treatment on skin bacterial dynamics and community ecology.

## INTRODUCTION

Skin represents a unique habitat, colonized by an equally unique set of microorganisms (1). Previous studies have analyzed these residents in-depth, describing a stable community distinguished by both inter- and intrapersonal differences (2, 3). This includes the skin’s ability to select for microbial residents at distinct biogeographic regions, each representing a niche with selective pressures that can influence cutaneous microbes (4). A number of studies have also tested the importance of these residents to human health, underscoring their ability to educate the immune system and protect against pathogenic skin microorganisms (5-8). Together, these studies have highlighted the importance of the skin microbiota, and outlined its role in host cutaneous defense.

Despite these findings, humans are constantly working to disrupt skin microbial communities in both clinical and non-clinical settings (9-12). While antimicrobial agents are largely employed to reduce infection by pathogenic microorganisms (13-15), these treatments can also act on resident cutaneous species (16-18). This is especially true for antiseptics, a group of antimicrobial agents used specifically for their indiscriminate mechanisms of action (19, 20). Antiseptics are a mainstay of modern medicine, but have also infiltrated daily routines in the form of gels, wipes, and sprays designed to sterilize the skin (21-23). As the significance of cutaneous resident microorganisms becomes increasingly apparent, assessing the impact of these treatments on the stability and resilience of skin microbiota becomes equally important. Indeed, we recently illustrated the potential for altered skin bacterial communities to impact colonization by *Staphylococcus aureus* in murine models, while others have expounded their importance in cutaneous diseases such as atopic dermatitis (24-26). These studies have highlighted the significance of skin microbial residents, and necessitated further research into treatment-derived perturbations.

To expand our knowledge in this regard, we designed a longitudinal treatment study to analyze how a “pulse” disturbance generated by topical antiseptics influences skin microbial community ecology using 16S ribosomal RNA (rRNA) gene sequencing. A single treatment was sufficient to elicit a significant impact on skin communities that was personalized and body site-specific. Certain microorganisms were more likely to be perturbed than others, with both abundance and bacterial identity representing key predictors of this response. These results further our understanding of stability and resilience of cutaneous microbial communities in the face of perturbation, and outline the potential for topical treatments to disrupt skin bacterial residents.

## RESULTS

Thirteen subjects, six females and seven males, were recruited to evaluate the effects of antiseptics on the skin microbiome. Treatments were applied to the volar forearm and the upper back to evaluate alternate skin microenvironments (dry and sebaceous, respectively), and each subject received identical treatments to control for interpersonal variability. Subjects received water and alcohol (80% ethanol) on contralateral body sites during their first series of visits, and povidone-iodine and chlorhexidine during their second series of visits. Swab specimens to analyze the microbiota were collected at baseline, prior to treatment, and at 6 time points post-treatment (1, 6, 12, 24, 36, and 72 hours; **Fig. S1a**) to assess longitudinal dynamics. Treated body sites were also accompanied by adjacent, untreated control sites, and visits were separated by at least two weeks to allow for microbial equilibration. Specific treatment topography, timing, and subject demographics are provided in **Fig. S1a** and **Table S1**. In total, 71,167,526 16S rRNA gene reads (hypervariable regions 1-3) were sequenced. Following quality control and filtering, the final study cohort represented 1,456 skin swab samples rarified to an even depth of 4,500 sequences per sample.

### Baseline characteristics of skin microbiota in study cohort

To validate our methods, we characterized the baseline communities of our study cohort. As previously reported (2, 4), we identified a strong impact of biogeography on the skin microbiota. Back communities were largely dominated by Propionibacteriaceae and Staphylococcaceae (**Fig. 1a**). By contrast, forearm communities were found to be more permissive, hosting increased proportions of additional taxa including Streptococcaceae and Corynebacteriaceae. Reflecting these community compositions, alpha diversity was significantly different between body sites, with the forearm exhibiting increased Shannon diversity, observed species, and equitability compared to the back (**Fig. 1b**). These metrics also highlighted the importance of interpersonal variability, as data points showed consistent grouping by individuals when assessing diversity at both body sites. When comparing these communities at the population-level, prominent clustering of subjects and body sites was observed by both weighted and unweighted UniFrac metrics (**Fig. 1c**). Comparisons of baseline communities also identified interpersonal variability and site-specificity as the most significant contributors to variation, followed by time and body symmetry respectively (**Fig. S1b-c**). These results confirm previous findings, and highlight the unique nature of resident skin bacterial communities.

**Fig. 1.**
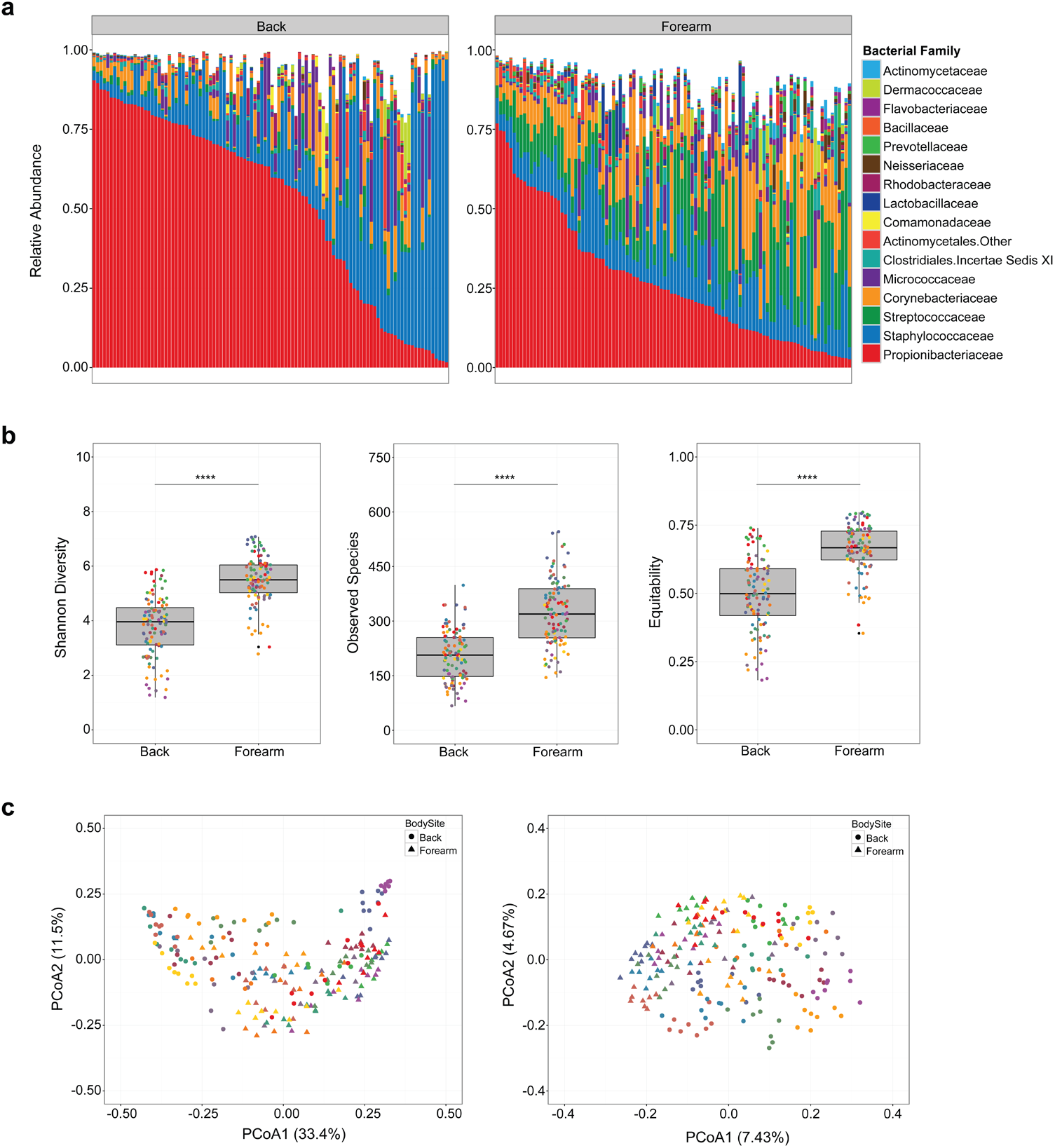
Skin bacterial communities exhibit site-specificity and interpersonal variability at baseline. (A) Family-level relative abundances of baseline communities for subjects at the forearm and back. Each bar represents an individual sample with eight samples per subject based on controls at adjacent and contralateral body sites for each visit series. (B) Alpha diversity of baseline communities at the forearm and back. Shannon diversity, observed species, and equitability are illustrated separately. Each point is colored by subject. (C) Weighted (left) and unweighted (right) UniFrac principal coordinates analyses of baseline samples. Each point is colored by subject and shaped by body site. **** P < 0.0001 by Wilcoxon rank sum test (Mann-Whitney U test).

### Treatment elicits subtle but personalized and site-specific shifts to skin bacterial community structure

We first compared baseline microbial communities to post-treatment communities at the 1 hour timepoint. Using weighted UniFrac and principal coordinates analysis, we observed minimal clustering of samples in response to treatment at the forearm and back, with none eliciting a significant shift in bacterial population structure (**Fig. 2a**). Because interpersonal differences were the strongest contributors to variability in baseline samples, and could thus mask more subtle effects of treatment, we also compared individual subjects’ posttreatment communities to their baseline controls. Using this method we detected a significant effect of both water and alcohol at the forearm, but not the back, for 6 hours post-treatment (**Fig. 2b**). While both treatments caused a more robust shift in forearm communities than that seen in adjacent controls, neither shifted bacterial communities to a state outside that of the broader study cohort (**Fig. 2c**). Comparisons of Shannon diversity and bacterial burden also confirmed these effects with alcohol eliciting significant decreases in diversity at the forearm, but not the back. However, water and alcohol were found to decrease overall bacterial load at each body site (**Fig. S2a, b**).

**Fig. 2.**
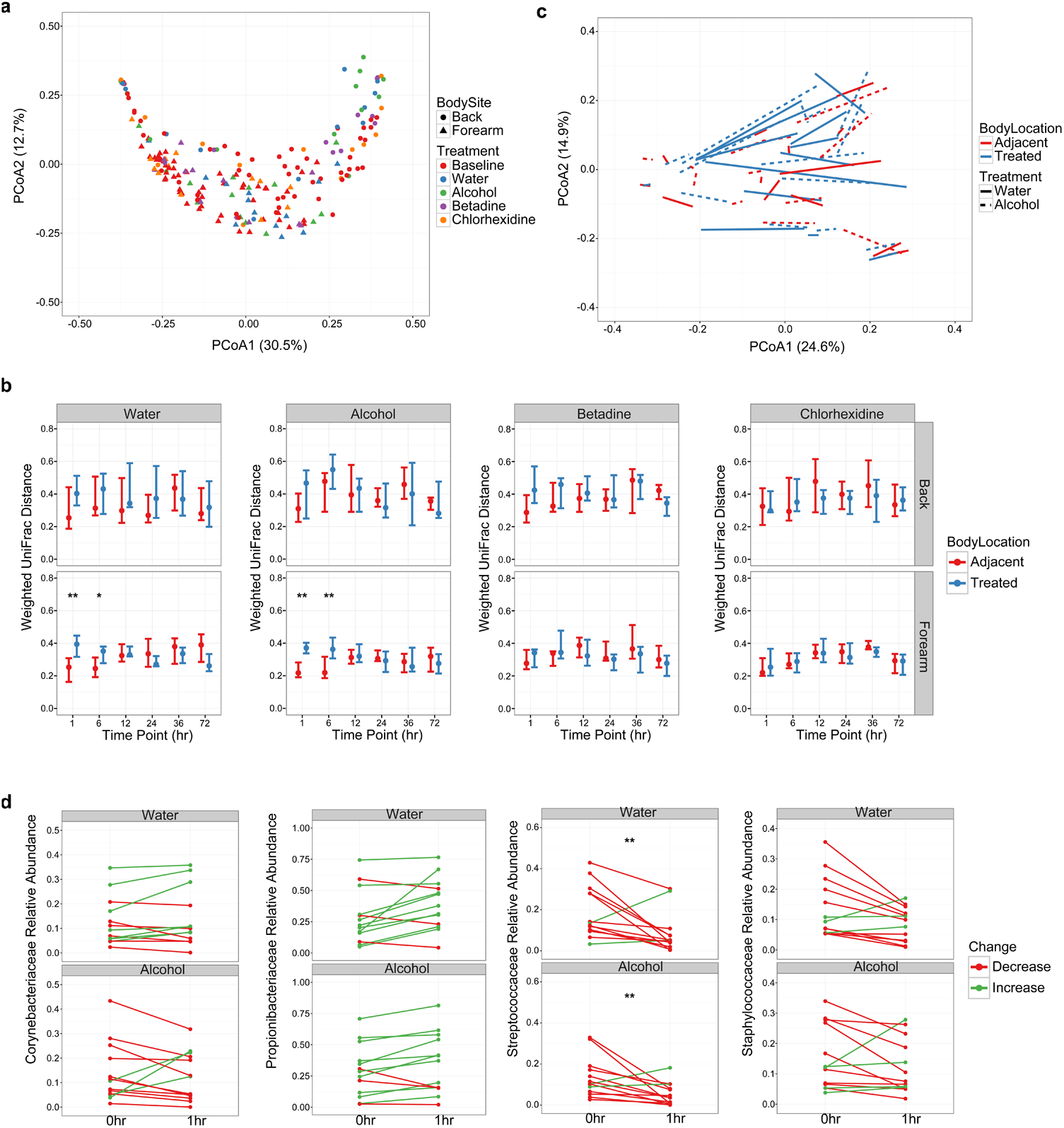
Treatment elicits personalized shifts in weighted comparisons of skin bacterial populations. (A) Principal coordinates analysis of weighted UniFrac distances for treated body sites at baseline and 1hr post-treatment. Each point represents a single sample, colored by treatment and shaped by body site. (B) Weighted UniFrac distances of subjects’ longitudinal time points compared to their individual baseline communities at treated and adjacent body sites. Points represent the median of participants. Error bars designate interquartile regions. (C) Subanalysis of weighted UniFrac distances visualized by principal coordinates analysis in subjects treated with water and alcohol at the forearm. Lines connect baseline and 1hr post-treatment samples for individual subjects, and line types designate treatment regimen. Line colors refer to treated body sites or their respective adjacent controls. (D) Comparison of relative abundances for the top 4 taxa at baseline and 1hr post-treatment with water or alcohol. Each line represents an individual subject colored by an increase or decrease in relative abundance following treatment. * P < 0.05, ** P < 0.01 by Wilcoxon rank sum test (Mann-Whitney U test).

To determine the taxa most responsible for these shifts, we focused our analyses on bacterial families with the greatest abundances prior to treatment. Corynebacteriaceae, Propionibacteriaceae, Streptococcaceae, and Staphylococcaceae were selected, as they represented a mean relative abundance of approximately 70% in pretreatment samples. Most taxa did not appear significantly altered despite variable changes in relative abundances of these taxa in our study cohort (**Fig. 2d**). Only Streptococcaceae was significantly decreased in response to treatment at the forearm, although both Propionibacteriaceae and Staphylococcaceae were also disrupted in nearly all subjects.

### Treatment depletes skin bacterial community membership and richness

We next investigated whether treatment could elicit more significant changes to skin bacterial community membership by unweighted metrics, which are agnostic to the relative proportions of bacterial taxa. In contrast to weighted comparisons, these tests revealed a prominent shift in bacterial communities following treatment at both the forearm and back (**Fig. 3a**). Moreover, when comparing treated communities to their baseline controls, both the back and forearm were found to be significantly disrupted by water, alcohol, and povidone-iodine compared to adjacent controls (**Fig. 3b**). To evaluate the underlying cause of this shift, we analyzed the effect of treatment on the total number of observed species. We found that changes to community membership were largely driven by a decrease in bacterial richness, with water, alcohol, and povidone-iodine all significantly reducing the number of observed species compared to adjacent controls (**Fig. S3a**).

**Fig. 3.**
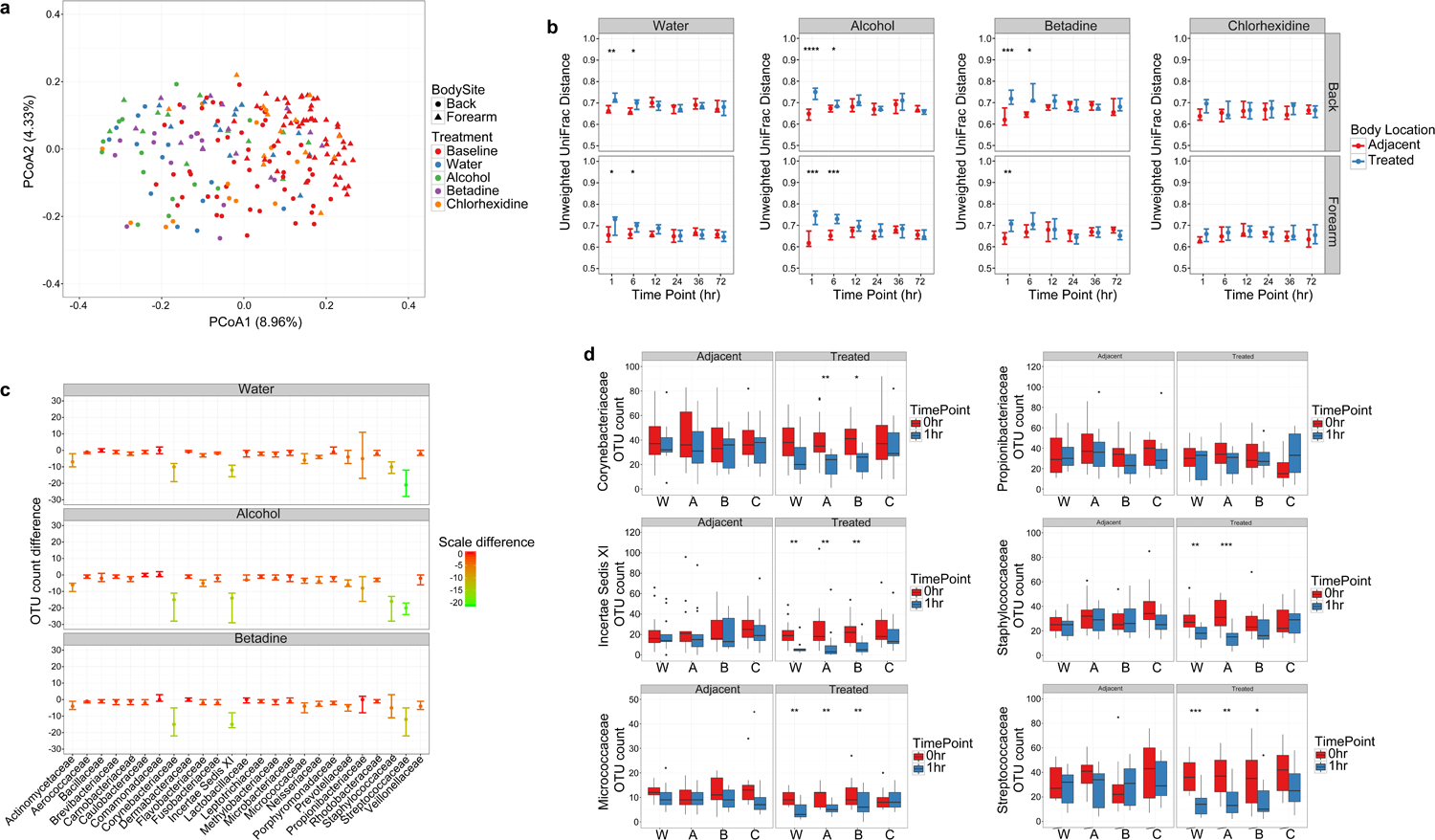
Treatment results in distinct alterations to skin bacterial residents by unweighted metrics. (A) Visualization of unweighted UniFrac distances by principal coordinates analysis for treated body sites at baseline and 1hr post-treatment. Each point represents a single sample, colored by treatment and shaped by body site. (B) Comparison of unweighted UniFrac distances for baseline and post-treatment communities in response to treatment at the forearm and back. Points represent the median of participants. Error bars designate interquartile regions. (C) Difference between OTU counts for the top 25 families at the forearm for baseline and 1hr post-treatment samples in response to water, alcohol, and povidone-iodine treatment. Points represent the median of participants and are colored by scaled differences in total count. Error bars designate interquartile regions. (D) Box and whisker plots of OTU counts for major taxa at adjacent and treated body sites of the forearm between baseline and 1hr time points. * P < 0.05, ** P < 0.01, *** P < 0.001, **** P < 0.0001 by Wilcoxon rank sum test (Mann-Whitney U test).

To further investigate these results, we also tested the effect of treatment on the membership of individual bacterial families. We found that Corynebacteriaceae, Incertae Sedis XI, Micrococcaceae, Staphylococcaceae, and Streptococcaceae were the most prominently disrupted taxa at both the forearm and back (**Fig. 3c; Fig. S3b**). Moreover, when comparing the richness of these taxa at treated and adjacent body sites, we found that each of these families were significantly decreased at treated, but not untreated, areas of the skin (**Fig. 3d; Fig. S3c**). Notably, this effect did not extend to all highly abundant families, as Propionibacteriaceae remained largely unchanged regardless of treatment or body site. This suggests that certain bacteria may be less susceptible than others when assessing treatment-derived alterations to bacterial membership, and underscores the differential ability of weighted and unweighted metrics to detect disruptions in bacterial populations.

### Chlorhexidine retains free bacterial DNA at the skin surface

Given its proven efficacy against pathogenic microorganisms in hospital settings (27), we were particularly struck that chlorhexidine did not elicit significant shifts in bacterial community membership or structure. As chlorhexidine is known to cause allergic and dermatologic irritation in a subset of individuals (28), we hypothesized that acute treatment results in changes to the skin barrier that allows for enhanced binding of free DNA released from dead bacteria. To test this hypothesis, we evaluated a subset of our subjects for alterations in skin barrier function by transepidermal water loss (TEWL) in response to treatment. We reasoned that if chlorhexidine were to alter the skin, making it more likely to bind free DNA, we should observe an increase in TEWL similar to that seen in patients with atopic dermatitis and other dermatologic conditions (29, 30). Upon testing, however, we found no significant differences in TEWL when comparing treatments to each other, or to baseline controls at 1hr and 6hr post-treatment (**Fig. S4a**).

We furthered hypothesized that chemical properties inherent to chlorhexidine were responsible for its lack of observed effect. To evaluate this question, we applied marker bacterial DNA to the skin of mouse dorsa, and tested its persistence following treatment with water, alcohol, povidone-iodine, or chlorhexidine. We observed that chlorhexidine uniquely retained free bacterial DNA, with the total amount of marker bacterial DNA exceeding that of other treatment regimens at 1hr post-treatment by over 10-fold on average (**Fig. S4a, b**). To test whether this effect could persist for multiple hours post-treatment, we also evaluated the quantity of DNA at 6hr post-treatment. Similar to 1hr time points, mice treated with chlorhexidine retained more DNA at the skin surface compared to other treatment regimens (**Fig. S4c**). These experiments suggest that failure to detect differences in resident skin microbiota following chlorhexidine treatment were likely due to a unique ability of this antiseptic to bind bacterial DNA to the skin surface, and not necessarily a deficiency in antibacterial activity. We therefore focused additional investigations on water, alcohol, and povidone-iodine treatments only.

### Treatment elicits convergence of skin microbiota at distinct community types

We next used an unsupervised approach to determine if a conserved microbial signature defined post-treatment microbial communities. Dirichlet multinomial mixture (DMM) modeling utilizes probability distributions to establish a prior of metacommunities (31). Clusters can then be generated based on the similarity of a sample to a given metacommunity. Using this approach with our study cohort, DMM models identified 8 distinct clusters at the forearm, with individual subjects often being dominated by a single community type (**Fig. S5a-b**).

Despite these interpersonal differences, however, we observed a prominent convergence to DMM cluster 1 in response to all treatments, an effect that was not observed at adjacent body sites (**Fig. 4a; Fig. S5c**). DMM cluster 1 was differentiated by decreased bacterial diversity, specifically richness (**Fig. 4b**). This particular cluster also displayed fewer taxon-specific attributes, suggesting a normalization of bacterial residents in response to treatment (**Fig. 4c**). In contrast to the forearm, back communities did not converge on a single community type following treatment (**Fig. S5d-e**). These data verify that treatment elicits reproducible changes to skin bacterial communities, namely depletion of community membership and diversity, but also underscores the importance of body site to calculations of resident stability.

**Fig. 4.**
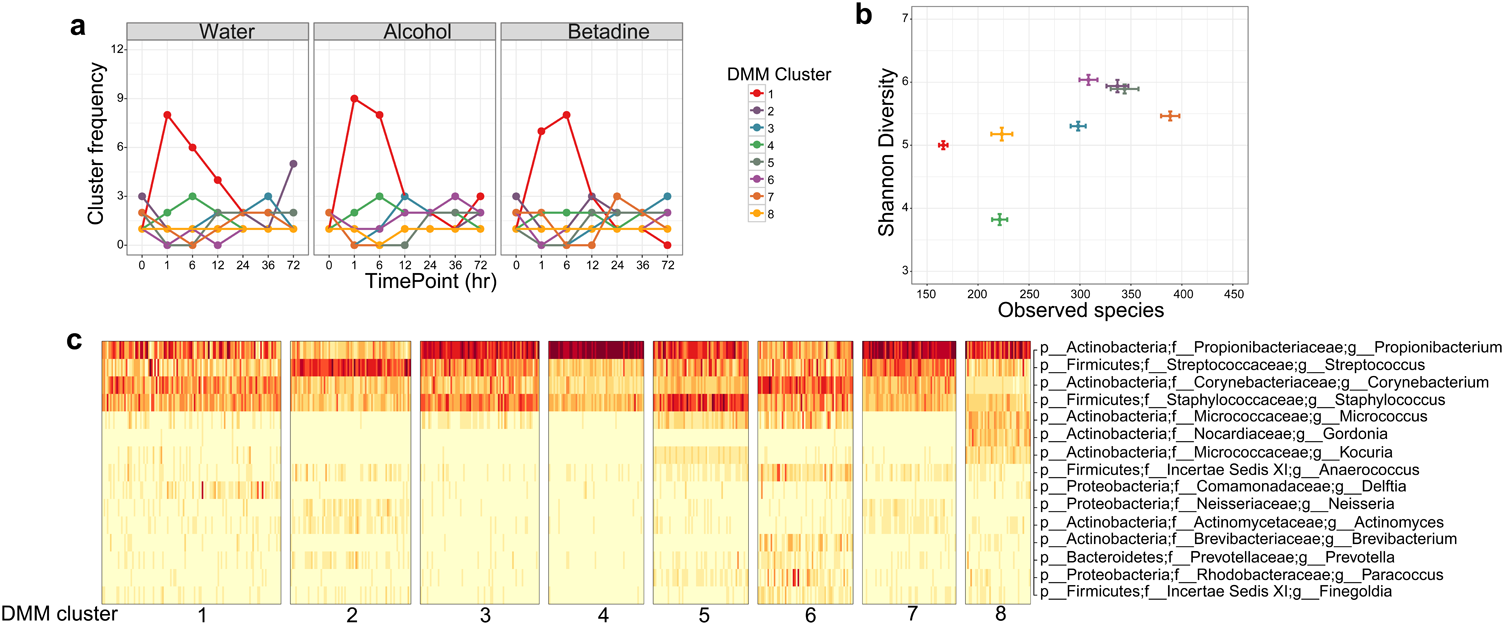
Dirichlet multinomial modeling identifies convergence at distinct forearm community types following treatment. (A) Longitudinal frequencies of DMM clusters in response to treatment with water, alcohol, and povidone-iodine. (B) Shannon diversity and observed species counts of individual DMM clusters. Data are presented as mean ± s.e.m. (C) Heat map of square root counts for the top bacterial taxa contributing to cluster identity. Dark bars correspond to greater counts.

### Highly abundant bacterial families are the greatest contributors to treatment-derived changes in skin bacterial communities

Our initial analyses suggested that certain bacterial taxa were disrupted more significantly by treatment than others. To assess this hypothesis, we tested characteristics shown to influence variation in untreated settings. We reasoned that the most variable taxa in the absence of treatment were also the most likely to be altered in response to topical intervention. As previous analyses have identified intermediately abundant taxa as the most susceptible to temporal fluctuation (32), we started by assessing the baseline variance of these taxa in our study cohort. Specifically, we compared the variance of bacterial residents at adjacent, contralateral, and temporally-controlled body sites to their mean relative abundances. Similar to previous findings, we observed a distinct second-order, power-law relationship in skin bacterial residents, with intermediately abundant members varying the most in untreated, baseline communities (**Fig. S6a**).

To test which taxa were specifically responsible for these shifts, we assessed baseline variance at the family level for each subject at the forearm and back. We found that Propionibacteriaceae, Streptococcaceae, Staphylococcaceae, Corynebacteriaceae, Micrococcaceae, and Incertae Sedis XI constituted the most variable groups in baseline communities (**Fig. S6b, c**). Interestingly, rather than representing only intermediately abundant taxa, however, these families were often the most abundant residents in our study cohort, and also the most likely to vary in response to treatment. To investigate this discrepancy within the literature more directly, we again compared the variance of baseline taxa in our study cohort to their mean relative abundances, but this time we further controlled for both inter-individual differences and body site-specificity. While we had previously observed a second-order relationship when aggregating subjects and body sites, stratification resulted in a more nuanced effect, with the variance of taxa frequently plateauing when plotted against their mean relative abundances (**Fig. 5a**). Indeed, top taxonomic groups often exhibited both the greatest levels of variance and the greatest mean relative abundances, especially in the case of Propionibacteriaceae. Together, these results suggest that intermediately abundant skin bacteria are the most likely to fluctuate at higher levels of comparison, but that predominant taxa are more variable when assessing personalized, biogeographic regions.

**Fig. 5.**
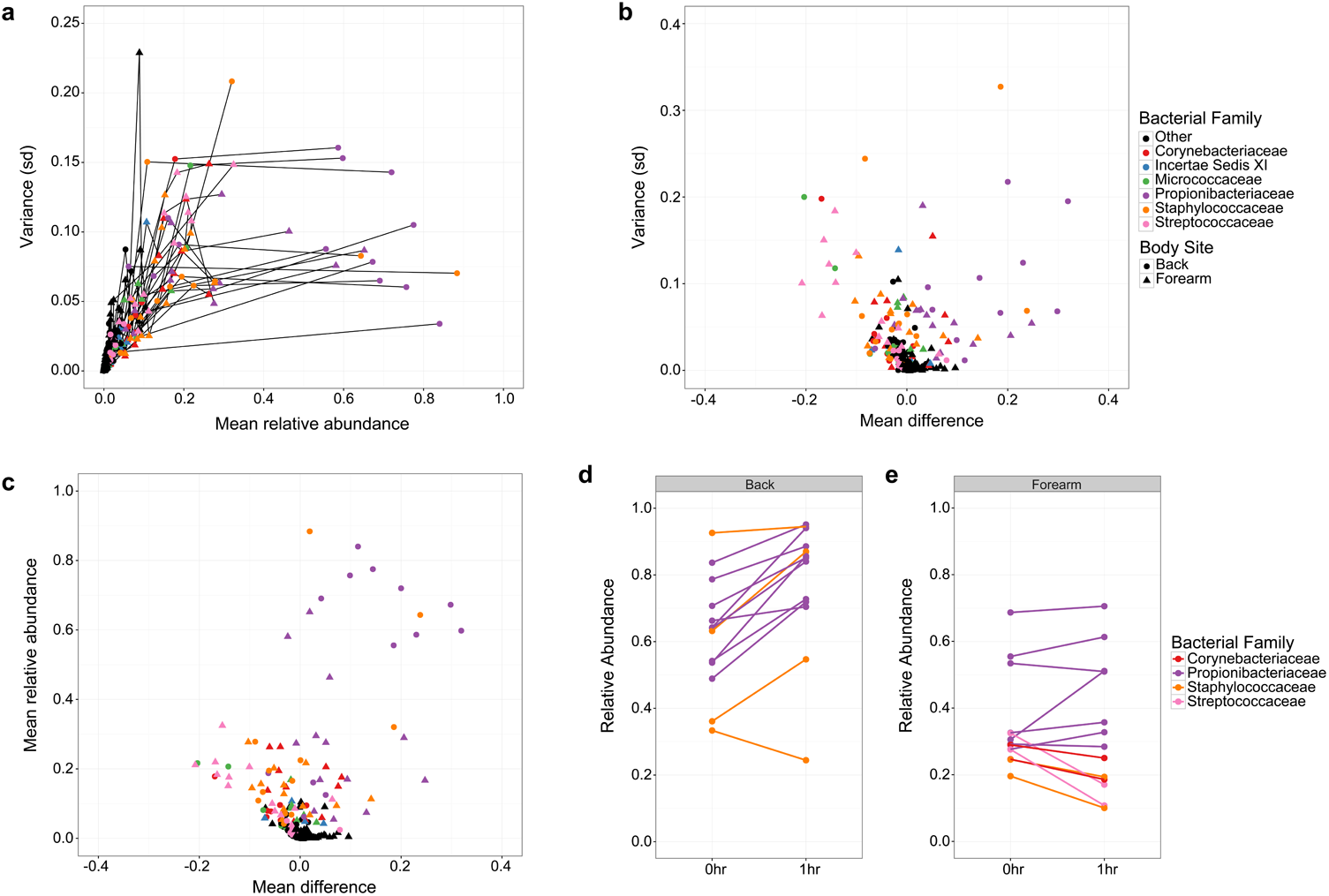
Baseline variance and abundance are indicators of treatment-derived alterations to the skin microbiota. (A) Family-level comparison of the baseline variances (standard deviation) and mean relative abundances for subjects at the forearm and back. Each point represents the values for bacterial families of an individual subject, shaped by body site and colored by family. Lines connect families of an individual subject and body site. “Other” designations refer to any bacterial family different from the listed members (B) Baseline variance of bacterial families plotted against their mean treatment effect in response to water, alcohol, and povidone-iodine treatment at the forearm and back. (C) Mean relative abundance of bacterial families at baseline compared to mean treatment effects at the forearm and back. (D, E) Mean difference in relative abundance of the most dominant taxon per subject following treatment at the back (D) and forearm (E). Each point represents a single subject colored by bacterial family identity.

We next tested whether taxonomic variation at baseline could be used as an indicator of post-treatment effects. Specifically, we compared the baseline variance of bacterial families to their mean response following water, alcohol, and povidone-iodine treatments. We found that the taxa most likely to vary in the absence of treatment were indeed the most likely to change in response to topical intervention, with decreases in the relative proportions of most taxa being offset by increases in Propionibacteriaceae (**Fig. 5b**). We observed that interpersonal variability was a strong contributor to this trend, as subjects with low variation of a given bacterial family were also less likely to exhibit shifts by those residents following treatment. This trend was recapitulated when comparing the mean relative abundances of taxa to their mean treatment response as well. Once again, the greatest differences were observed within the Propionibacteriaceae family, which was both the most abundant bacterial family and the most likely to increase following treatment (**Fig. 5c**). In all, these results indicate that both abundance and variation in untreated controls can inform treatment-derived effects, but that bacterial identity is also an important variable when measuring overall community response.

### Body site specificity informs fluctuations of the most abundant bacterial taxa

During these analyses, we noted that Propionibacteriaceae, unlike other taxa, often increased in relative abundance following treatment of the back. We also found that a subset of subjects exhibited similar dynamics when Staphylococcaceae was their most abundant taxon. Because we observed a decrease in bacterial load following treatment, these increases in relative proportions were unlikely to represent increases in absolute abundance. However, they did suggest a personalized response in which the most abundant taxon per subject was also the most likely to persist following treatment. To test this hypothesis, we compared the levels of each subject’s most abundant taxon at baseline to its mean relative abundance following water, alcohol, and povidone-iodine treatment. We found that in all cases but one, the most abundant taxon at the back increased in relative proportion following treatment regardless of identity, indicating a distinct competitive advantage (**Fig. 5d**).

Unlike the back, only three subjects had taxa at the forearm with > 50% relative abundance. However, we still observed an increase in the relative proportions of Propionibacteriaceae in multiple subjects following treatment, although this effect was not absolute (**Fig. 5e**). This trend did not extend to all skin residents, as Corynebacteriaceae, Staphylococcaceae, and Streptococcocaceae all decreased in abundance at the forearm, regardless of status. These results thus verify that abundance can be used to predict treatment effects, but also highlights the importance of body site to these particular outcomes.

### Lowly abundant members of predominant bacterial families are the most likely to vary in response to treatment

Because our previous investigations outlined the importance of abundance and bacterial identity to treatment-derived alterations, we further hypothesized that relative abundance could be used to predict the fluctuations of all taxa. To test this, we partitioned OTUs into highly or lowly abundant groups based on an abundance threshold of 0.5% - a value chosen from the inflection point of OTU counts at baseline (**Fig. S7a**). We observed a significant decrease in the number of lowly abundant OTUs following treatment at both the forearm and back (**Fig. 6a**), an effect largely due to decreases in Corynebacteriaceae, Incertae Sedis XI, Staphylococcaceae, and Streptococcaceae (**Fig. 6b-c; Fig. S7b-c**). By contrast, when evaluating highly abundant OTUs, only Streptococcaceae at the forearm and Corynebacteriaceae at the back were significantly reduced, a result which did not significantly decrease the total number of highly abundant OTUs. Similar to previous results, we also observed no significant differences in the membership of Propionibacteriaceae, regardless of abundance or body site. These findings confirm that bacterial identity represents a critical factor when evaluating skin resident stability, and underscores the importance of abundance to predictions of treatment response.

**Fig. 6.**
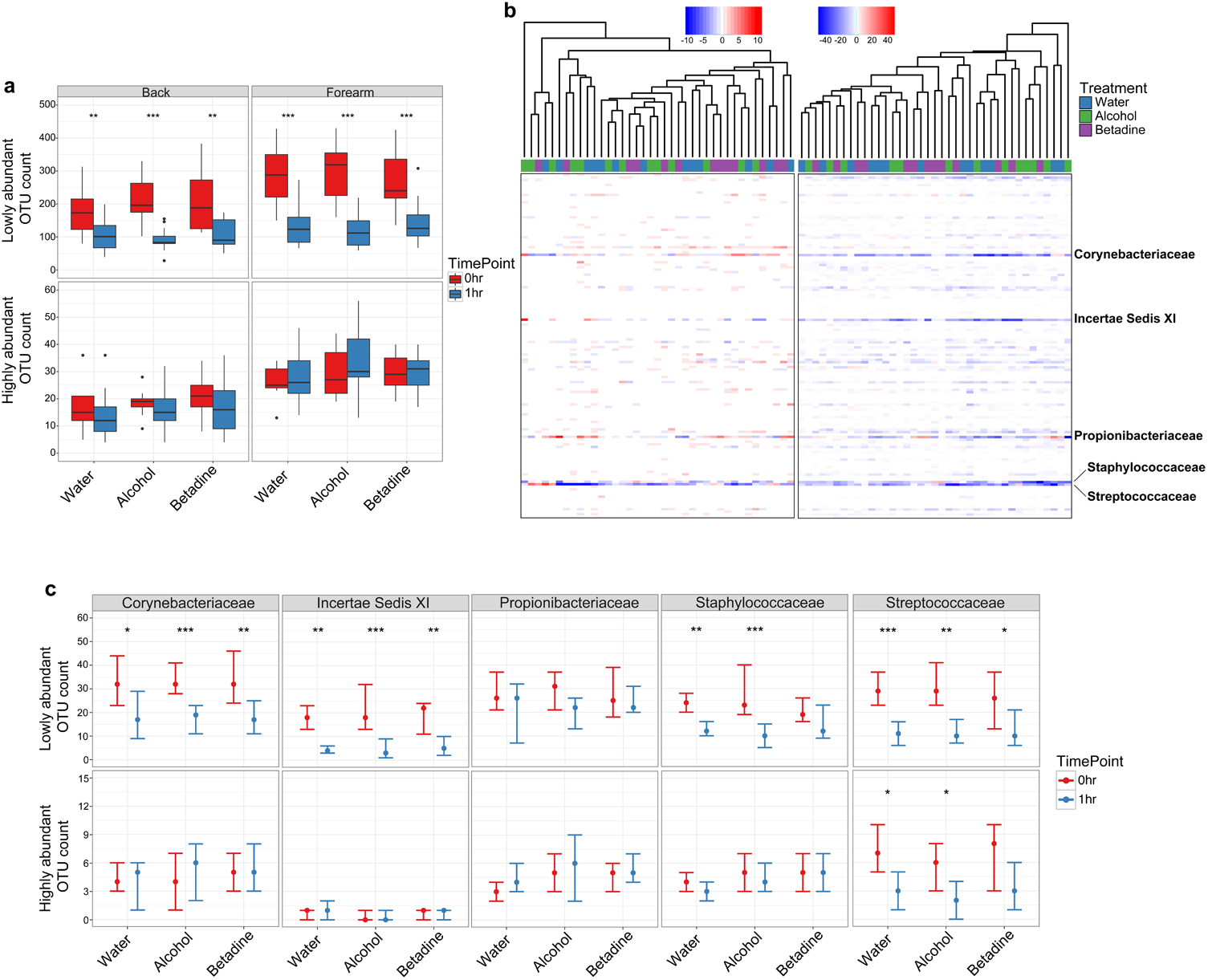
Lowly abundant members of prominent taxa are the greatest contributors to treatment effects at the skin surface. (A) Box and whisker plots of lowly and highly abundant OTU counts as defined by a 0.5% relative abundance threshold following treatment at the forearm and back. (B) Heat map of differences in forearm OTU counts between baseline and 1hr post treatment with water and antiseptics. Each column represents the difference measured for a single subject and treatment, and each row represents a bacterial family. Samples are clustered by the Unweighted Pair Group Method with Arithmetic means (UPGMA). Color-coded bars above the graph designate treatments for each sample. (C) Comparison of lowly and highly abundant OTU counts at the forearm in major taxonomic families at baseline and 1hr post-treatment. Points represent the median of the study cohort. Error bars designate interquartile regions. * P < 0.05, ** P < 0.01, *** P < 0.001 by Wilcoxon rank sum test (Mann-Whitney U test).

## DISCUSSION

The skin microbiota has proven essential to numerous functions in cutaneous health and disease (5-8). However, few studies have assessed the impact of disruption on these communities, or their dynamics following antimicrobial stress. Herein, we present the impact of topical antiseptics on human skin bacterial populations, and outline the importance of key variables to this response.

When evaluating treatments at a comparative level, we found water, alcohol, and povidone-iodine to have similar effects on skin bacterial residents. These results underscore the generalized nature of topical interventions to reduce inhabitance by mechanical cleansing (19). This result has been well-established in culture-based systems, where reports have outlined the potential for certain topical treatments to both kill, and remove, pathogenic microorganisms, with each feature playing an important role in infection control (33, 34). Mild, non-antibacterial soaps are also used with the sole purpose of clearance, further emphasizing the importance of this mechanism to skin hygiene and community disruption (35, 36).

Unlike alcohol and povidone-iodine, chlorhexidine was found to elicit only minor shifts in skin bacterial residence, which was surprising given the ability of chlorhexidine to reduce infections in clinical settings (37). We found that chlorhexidine was uniquely proficient at retaining bacterial DNA on the skin for up to 6hr post-treatment. Further research is necessary to elucidate the mechanism by which chlorhexidine achieves this feat. It is interesting to note that chlorhexidine is a cationic molecule, distinguished by an ability to bind the epidermal surface for multiple hours post-treatment (38). As such, the potential exists that this antiseptic may retain bacterial DNA through a dual interaction with skin keratinocytes and negatively charged DNA backbones.

While no study to date has investigated the impact of antiseptics on human skin microbiota by sequencing, others have assessed the effects of hand-sanitizers and soaps (39, 40). These studies have largely supported culture-based tests, outlining the importance of conserved mechanisms to topical treatment response. For example, a recent study by Zapka, et al. found that water and hand washing often elicited similar alterations to the skin microbiota as alcohol-based hand sanitizers (39). A recent comparison of mild and antibacterial soaps has further confirmed these results, showing minimal differences when comparing their impact on colonizing levels of *S. epidermidis* (40).

Like the abovementioned studies, our initial analyses suggested a relatively minor impact of treatment on resident microbiota. However, after controlling for personalization and body site-specificity we observed the true impact of our treatment regimens on community diversity and resilience, including the finding that treatment elicited the strongest effects in low-level skin inhabitants. Highly abundant species likely exist at a given skin niche due to an ability to resist acute host-derived and external stressors. As the skin is often colonized by particular strains with temporal stability for years in length (32, 41), this outlines a system by which multiple taxa may exist on the skin surface at a given time, while only a subset are uniquely adapted for long-term colonization. These observations pair well with previous, culture-based analyses where highly abundant skin residents have been shown to persist in response to various treatments while transient, low-level bacteria represent a less stable group of community inhabitants (42-44).

We also found that bacterial identity could influence treatment response, with predominant skin taxa often more significantly disrupted than other residents. This finding underscores the ecological advantages seen in bacterial families such as Propionibacteriaceae, Staphylococcocaceae, Streptococcaceae, and Corynebacteriaceae. The prevalence of these taxa at baseline in most subjects likely illustrates their ability to utilize conserved resources at and within the skin surface (45). Upon the introduction of treatment-derived stressors, however, a generalized selective advantage is no longer enough, leading to the persistence of only the most resilient and well-adapted members of these groups.

Treatment-derived alterations were also found to be dependent upon body site, with the back representing a more stable habitat than the forearm in most tests. Notwithstanding, both body sites were susceptible to a loss of lowly abundant OTUs in many predominant skin residents, emphasizing the reproducibility of this particular observation. This result did not extend to all major taxa, as members of the Propionibacteriaceae family persisted regardless of body site. We believe this particular effect could be due to an inherent resilience of Propionibacteriaceae, or its increased abundance at deeper, newly exposed layers of the skin. Regardless, the persistence of this taxon likely represents a unique opportunity to thrive in the post-treatment setting.

In all, this study furthers our understanding of skin bacterial dynamics and elucidates the effects of topical treatments on cutaneous resident populations. While we observed a similar impact of water and the antiseptics alcohol and povidone-iodine on skin inhabitants, we note that our studies were designed to assess the totality of skin residents in healthy individuals. As such, we caution against the application of these findings to clinical settings in which the dynamics of pathogens and commensals are highly skewed. Indeed, previous studies have described, in-depth, the utility of antiseptics in these particular environments (46-48). As our study assesses only the effect of acute stressors, or a “pulse” disturbance, we also advocate for further research into long-term treatment regimens more characteristic of a “press” disturbance. The potential exists that more lasting perturbations may elicit even greater shifts to human skin bacterial communities, an important consideration when evaluating the nexus of host-microbial interactions.

## MATERIALS & METHODS

### Human subjects and sample collection

All protocols were approved by the Institutional Review Board of the University of Pennsylvania, and written informed consent was obtained for all study participants prior to sampling. Thirteen healthy subjects aged 23-30 (median:27, 6 females) and without chronic skin disorders were recruited to participate in a controlled skin antiseptic study (**Table S1**). Subjects were required to be >21 years of age, and free of oral or topical antibiotics within 6 months of their first visit. Subjects were asked to 1) refrain from showering for 24 hours prior to the first visit and until after their 36 hour visit, and 2) refrain from use of soaps or topical products containing antimicrobials 1 week prior to sampling and during the entire study. Demographic data were collected as well as usage of topical products, medications, personal care routines. Subjects were swabbed at baseline, using a Catch-All Collection swab (Epicentre, discontinued) moistened in water (UltraPure Distilled Water, Invitrogen) and then administered one of four treatments for 1.5 minutes. Each participant received water (UltraPure Distilled Water, Invitrogen) and alcohol (80% ethanol) on contralateral forearm or back body sites during their first visit series, and povidone-iodine (10%; Betadine), and chlorhexidine (chlorhexidine-gluconate 4%) during their second visit series (**Fig. S1a**). Visit series were separated by at least two weeks to allow for microbial equilibration. Following treatment, subjects were swabbed at 1hr, 6hr, 12hr, 24hr, 36hr, and 72hr post-treatment at both treated and adjacent body sites. Swabbed regions were delineated by a skin marker to ensure that the same body site was swabbed at longitudinal time points. Subjects were instructed to refrain from showering for at least 12 hours prior to each time point.

### Transepidermal water loss

Transepidermal water loss was measured in a subset of four subjects using a Tewameter TM300 (Courage+Khazaka, Cologne, Germany) according to the manufacturer’s instructions. Briefly, subjects were equilibrated for at least 10 minutes prior to testing. Noninvasive probes were then pressed to the skin at baseline, 1hr, and 6hr post-treatment with water, alcohol, povidone-iodine, or chlorhexidine to measure changes in skin epidermal barrier function. Each process was repeated at both the forearm and back to assess differences by body site.

### Bacterial DNA isolation, 16S rRNA gene sequencing, and qPCR

Bacterial DNA was extracted as described previously (49) using the Invitrogen PureLink kit. PCR and sequencing of the V1V3 hypervariable region was performed using 300-bp paired end chemistry and barcoded primers (27F, 534R) on the Illumina MiSeq platform. Accuprime High Fidelity Taq polymerase was used for PCR cycling conditions: 94 °C for 3 min; 35 cycles of 94 °C for 45 sec, 50 °C for 60 sec, 72 °C for 90 sec; 72 °C for 10 min. For bacterial load comparisons, 16S rRNA genes were amplified by qPCR using Fast SYBR Green Master Mix (Fisher Scientific) and the optimized primers 533F, 902R. Samples were compared to standard curves generated from known concentrations of serially diluted bacterial DNA to calculate burden.

### Microbiome analysis

The datasets generated and analyzed during the current study are available in the NCBI Short Read Archive under BioProject: PRJNA395539. Sequences were preprocessed and quality filtered prior to analysis, and QIIME 1.7.0 was used for microbiome evaluation (50). Briefly, sequences were *de novo* clustered into OTUs based on 97% similarity by UClust (51), and taxonomy was assigned to the most abundant representative sequence per cluster using the RDP classifier (52). Sequences were aligned by PyNAST (53), and chimeric sequences were removed using ChimeraSlayer (54). Sequences with calls to Unclassified, Bacteria;Other, or Cyanobacteria were removed in addition to singletons. Antiseptics and negative controls were similarly sequenced and analyzed for possible contaminating sequences, with no OTUs being found at consistently high levels. All samples were rarified to 4,500 sequences, and samples below this cut-off were removed from downstream analyses. Alpha and beta diversity matrices and taxonomy tables were formulated in QIIME. Statistical analysis and visualization were performed in the R statistical computing environment (55).

### Dirichlet multinomial mixture models

Subsampled OTU counts were aggregated at the highest level of taxonomic classification. Samples were separated by body site and spurious taxa in less than 1% of samples were removed. Clusters were generated separately on forearm and back samples using the R package Dirichlet Multinomial (v1.14.0), and community types for each body site were calculated based on absolute minima from Dirichlet components and Laplace approximations of model evidence (31). Samples were assigned to final community types based on posterior probabilities.

### DNA retention

C57BL/6J mice were bred and maintained in specific pathogen free conditions at the University of Pennsylvania. All animal protocols were reviewed and approved by the University of Pennsylvania Institutional Animal Care and Use Committee. Eight to fifteen-week-old males and females were randomized to control for differences in age and gender, and each mouse was housed singly to avoid cross-contamination. Mice were shaved at the dorsum and acclimated for at least 2 days prior to experimentation. 5-6 ng/ul of extracted *Escherichia coli* DNA was applied to mouse dorsa and permitted to dry for 1hr prior to treatment. Mice were then administered water, alcohol, povidone-iodine, or chlorhexidine for 1.5 minutes, similar to human experiments, and swabbed at 1hr and 6hr post-treatment. DNA was extracted using the Invitrogen PureLink kit, and *E. coli*-specific DNA was amplified using qPCR primers to the *ycct* gene (56). Samples were compared to standard curves generated from known amounts of serially diluted *E. coli* DNA to calculate marker DNA concentrations.

## ACKNOWLEDGEMENTS

We thank Penn Next Generation Sequencing Core for sequencing support, the Penn Medicine Academic Computing Services for computing support, and members of the Grice laboratory for their underlying contributions. Funding for this work was provided by the NIH, National Institute of Arthritis, Musculoskeletal, and Skin Diseases (R00AR060873 and R01AR066663 to EAG) and the National Institute of Nursing Research (R01NR015639 to EAG). AJS is supported by a Department of Defense National Defense Science and Engineering Graduate fellowship. JSM is supported by NIH T32HG00046, Computational Genomics Training Grant. The content is solely the responsibility of the authors and does not necessarily represent the official views of the National Institutes of Health or the Department of Defense.

---

**Fig. S1.** Treatment regimen details and baseline community comparisons. (A) Diagram of sampling and treatment schedule for antiseptic study cohort. (B, C) Heat map of significances for weighted (B) and unweighted (C) UniFrac comparisons at the forearm and back for interpersonal, adjacent, contralateral, shortterm (1hr), and long-term (visit) baseline community samplings.

**Fig. S2.** Alpha diversity and bacterial load are decreased in response to certain treatment regimens. (A) Longitudinal comparisons of Shannon diversity for bacterial communities at adjacent and treated body sites of the back and forearm. (B) Bacterial load at the forearm and back for treated and adjacent body sites over time. Data is presented by median points and interquartile regions. * P < 0.05, ** P < 0.01 by Wilcoxon rank sum test (Mann-Whitney U test).

**Fig. S3.** Treatment elicits decreases in bacterial richness. (A) Longitudinal measurements of observed species for adjacent and treated body sites at the back and forearm. Data is presented by median points and interquartile regions. (B) Difference between OTU counts for the top 25 families at the back for baseline and 1hr post-treatment samples in response to water, alcohol, and povidone-iodine treatment. Points represent the median of participants and are colored by the scaled difference in total count. Error bars designate interquartile regions. (C) Box and whisker plots of OTU richness at the back for major taxa at adjacent and treated body sites between baseline and 1hr time points. * P < 0.05, ** P < 0.01, *** P < 0.001 by Wilcoxon rank sum test (Mann-Whitney U test).

**Fig. S4.** Effect of chlorhexidine on skin integrity and bacterial DNA retention. (A) Transepidermal water loss (TEWL) of subjects at the back and forearm in response to treatment with water, alcohol, povidone-iodine, and chlorhexidine. Each point represents an individual subject. Black bars denote median. (B) Concentration of marker bacterial DNA at baseline and 1hr and 6hr post-treatment. Each point represents an individual mouse. Black bars denote median. Baseline refers to background concentrations of marker DNA prior to testing.

**Fig. S5.** Dirichlet multinomial mixture (DMM) model analysis for bacterial communities at the forearm and back. (A) Frequency of forearm DMM clusters by subject. (B) Laplace approximations for Dirichlet components of forearm communities. Global minimum is represented with a red point. (C) Frequencies of forearm DMM clusters at adjacent body sites over time. (D) Laplace approximations for Dirichlet components of back communities. (E) Longitudinal frequencies of DMM clusters at the back for adjacent and treated body sites. (F) Shannon diversity and observed species counts of individual DMM clusters at the back. Data are presented as mean ± s.e.m.

**Fig. S6.** Variance of bacterial taxa at baseline. (A) Relationship between the mean relative abundances of bacterial families at baseline and their variance as measured by standard deviation. Each point represents a different bacterial family in an individual subject, shaped by body site. Data was fitted with a second-order curve to approximate taxonomic distributions. (B, C) Baseline variance of top 25 bacterial families at the back (B) and forearm (C). Points are colored by subject, and shaped by body site. Black bars represent median variance.

**Fig. S7.** Contribution of abundance to treatment-derived alterations in bacterial membership. (A) Sorted OTU abundances of all bacterial members in study cohort. Dashed red line represents 0.5% abundance threshold used to separate highly and lowly abundant OTUs. (B) Heat map of differences in bacterial family membership at the back between baseline and 1hr in subjects following treatment with water, alcohol, and povidone-iodine. Each column represents the sample of an individual subject, and each row represents a bacterial family. Samples are clustered by the Unweighted Pair Group Method with Arithmetic means (UPGMA). Color-coded bars above the graph designate treatments for each sample. (C) Comparison of lowly and highly abundant OTU counts at the back in major taxonomic families at baseline and 1hr post-treatment. Data is presented by median points and interquartile regions. * P < 0.05, ** P < 0.01, *** P < 0.001 by Wilcoxon rank sum test (Mann-Whitney U test).

